# Logical design of oral glucose ingestion pattern minimizing blood glucose in humans

**DOI:** 10.1101/352955

**Authors:** Masashi Fujii, Yohei Murakami, Yasuaki Karasawa, Yohei Sumitomo, Suguru Fujita, Masanori Koyama, Shinsuke Uda, Hiroyuki Kubota, Hiroshi Inoue, Katsumi Konishi, Shigeyuki Oba, Shin Ishii, Shinya Kuroda

## Abstract

Excessive increase in blood glucose level after eating increases the risk of macroangiopathy, and a method for not increasing the postprandial blood glucose level is desired. However, a logical design method of the dietary ingestion pattern controlling the postprandial blood glucose 2 level has not yet been established. We constructed a mathematical model of blood glucose control by oral glucose ingestion in 3 healthy human subjects, used the model to predict an optimal glucose ingestion pattern, and showed that the optimal ingestion pattern minimized the peak value of blood glucose level. Subjects orally ingested 3 doses of glucose by bolus or over 2 hours, and blood glucose, insulin, C-peptide and incretins were measured for 4 hours. We constructed an ordinary differential equation model that reproduced the time course data of the blood glucose and blood hormone levels. Using the model, we predicted that intermittent ingestion 30 minutes apart was the optimal glucose ingestion patterns that minimized the peak value of blood glucose level. We confirmed with subjects that this intermittent pattern decreased the peak value of blood glucose level. This approach could be applied to design optimal dietary ingestion patterns.

**In Brief:** As a forward problem, we measured blood glucose and hormones in three human subjects after oral glucose ingestion and constructed a mathematical model of blood glucose control. As an inverse problem, we used the model to predict the optimal oral glucose ingestion pattern that minimized the peak value of blood glucose level, and validated the pattern with the subjects.

**Highlights:**

- Modeling blood glucose concentrations predicts an intermittent ingestion pattern is optimal
- Human validation shows ingestion at 30-minute intervals limits peak blood glucose
- We provide a strategy to design optimal dietary ingestion patterns

## INTRODUCTION

In healthy people, blood glucose levels are stably maintained and show only a slight postprandial increase (Abdul-Ghani et al., 2006). However, massive postprandial increases in blood glucose levels emerge in patients with the type 2 diabetic mellitus (T2DM) and impaired glucose tolerance (Edelstein et al., 1997). This postprandial hyperglycemia requires prevention and treatment, because it is associated with increased risk of cardiac and cerebrovascular complications (Nakagami and DECODA Study Group, 2004). Postprandial blood glucose originates from dietary carbohydrates (Cahill, 2006). Some approaches to prevent postprandial hyperglycemia have thus far been reduction of dietary carbohydrate content, a change in the type of dietary carbohydrates, and ingestion of dietary fiber with meals (Schulze et al., 2004). However, the ideal type of pattern of ingestion of carbohydrate that minimize postprandial hyperglycemia is unknown.

Insulin, secreted from the pancreatic β cells, performs a pivotal role in homeostatic regulation of blood glucose levels. Insulin acts on the target organs such as muscle and liver, to promote uptake of glucose from the blood and suppress hepatic glucose production. Consequently, insulin decreases blood glucose levels and promotes the rapid recovery of increase in postprandial blood glucose. As blood glucose levels decrease, insulin secretion also decreases. Thus, the blood glucose level is maintained within a narrow normal range by the feedback relationship between blood glucose and insulin (Castillo et al., 1994).

Although insulin secretion is regulated mainly by blood glucose, it is also regulated by a family of circulating hormones called incretins (Seino et al., 2010). Incretins are hormones secreted from the gastrointestinal tract upon food ingestion, these hormones act on pancreatic β cells to promote insulin secretion. Gastric inhibitory polypeptide (GIP) and glucagon-like peptide-1 (GLP-1) are incretins (Fujimoto et al., 2009; Preitner et al., 2004; Seino et al., 2010; Vollmer et al., 2008). GIP is secreted from K cells of the upper small intestine (Inagaki et al., 1989; Takeda et al., 1987); GLP-1 is secreted from L cells of the lower small intestine (Bell et al., 1983; Orskov et al., 1993). Orally ingested glucose promotes incretin secretion into the small intestine, where it is absorbed and enters the blood. Blood glucose and incretin act cooperatively on pancreatic β cells to promote insulin secretion and increase circulating insulin levels (Parkes et al., 2001).

Postprandial hyperglycemia is identified with an oral glucose tolerance test (OGTT), in which a subject’s ability to tolerate a glucose load (glucose tolerance) is evaluated by measuring blood glucose level after an overnight fast and again 2 h after a 75-g oral glucose load (Stumvoll et al., 2000). Using time course data of glucose and insulin in the blood during the OGTT, many mathematical models have quantitatively evaluated the relationship between the blood glucose and insulin in humans (Bergman et al., 1979; Brubaker et al., 2007; Dalla Man et al., 2016, 2013, 2007, 2006; De Gaetano et al., 2013; Hill et al., 1997; Kabul et al., 2015; Kim et al., 2014; Overgaard et al., 2006; Pedersen et al., 2011; Riz et al., 2014; Røge et al., 2017; Salinari et al., 2011; Tura et al., 2001). These models consist of blood glucose and insulin, but not incretins (Bergman et al., 1979; Dalla Man et al., 2007, 2006; De Gaetano et al., 2013; Tura et al., 2001). Other mathematical models incorporate the incretins (Brubaker et al., 2007; Dalla Man et al., 2016; Kabul et al., 2015; Kim et al., 2014; Pedersen et al., 2011). In some models, blood glucose and incretin act independently on insulin secretion during the OGTT (Brubaker et al., 2007; Kabul et al., 2015; Kim et al., 2014); in others, blood glucose and incretin act cooperatively (Dalla Man et al., 2016; Pedersen et al., 2011). The effective action of incretins on the insulin secretion in mathematical models remains to be determined.

One application of mathematical models is the ability to make prediction. Published mathematical models of blood glucose and insulin have been used to predict blood glucose levels after glucose administration. We require a solution of a pair of forward and inverse problems to obtain an optimal design of input pattern. Firstly, we need a dynamics model to predict the temporal pattern as a consequence of a given input pattern. This mode of prediction is a forward problem: The prediction is an “output pattern” related to the input pattern. Secondly, optimal input pattern should be determined so as to minimize the outcome that is defined as an arbitrarily given objective function of the predicted output pattern. This mode of prediction is an inverse problem: The prediction is an “input pattern” that produces an optimal output pattern. There are many established methods that use complex ordinary differential equations to solve the forward problem of predicting output patterns, but few methods exist to solve the inverse problem of predicting input patterns. Recently, we proposed a mathematical framework to estimate an input pattern that produces a defined output pattern (Murakami et al., 2017).

Here, we constructed mathematical models with either glucose-independent and/or glucose-cooperative roles of incretins on insulin secretion. We used the models to predict an optimal glucose ingestion pattern that controls blood glucose level. Because blood glucose level is the output pattern, this represents using the model to solve an inverse problem. We measured blood glucose, insulin, GIP and GLP-1 before and after oral glucose ingestion with different doses and ingestion durations for three subjects. As a forward problem, we constructed a mathematical model of blood glucose (output) in response to orally ingested glucose (input) for each subject. As an inverse problem, we optimally designed glucose ingestion pattern that minimized the peak value of blood glucose level for each subject. Each subject had an optimized pattern of ingestion that was intermittent. We validated blood glucose level by the predicted intermittent ingestion pattern for each subject and found that the intermittent ingestion pattern decreased the peak value of blood glucose level compared with the blood glucose levels that occurred with bolus or 1 h-continuous ingestion patterns. Thus, we provide the logical design of oral glucose ingestion pattern that minimizes the peak value of blood glucose level in humans, using an approach of combination of a forward and an inverse problems, which can be widely applied to design optimal dietary ingestion patterns for human health.

## RESULTS

### Measurement of Blood Glucose and Blood Hormones Before and After Oral Glucose Ingestion

To obtain the data for developing the model, we monitored the effect of ingestion of different amounts of glucose in different temporal patterns of ingestion on blood glucose and hormone levels (Figure 1). In 6 separate experiments, the three healthy volunteers either rapidly consumed one of three doses of glucose (25 g, 50 g, 75 g) or consumed the glucose over 2 hours (see STAR Methods A.1, A.2). The rapid ingestion paradigm is referred to as bolus ingestion and the slow ingestion paradigm as 2 h-continuous ingestion. Prior to glucose ingestion and after glucose ingestion, we measured levels of blood glucose, insulin, C-peptide, intact GIP (designated GIP hereafter), and intact GLP-1 (designated GLP-1 hereafter) (see STAR Methods A.2).

**Figure 1.**
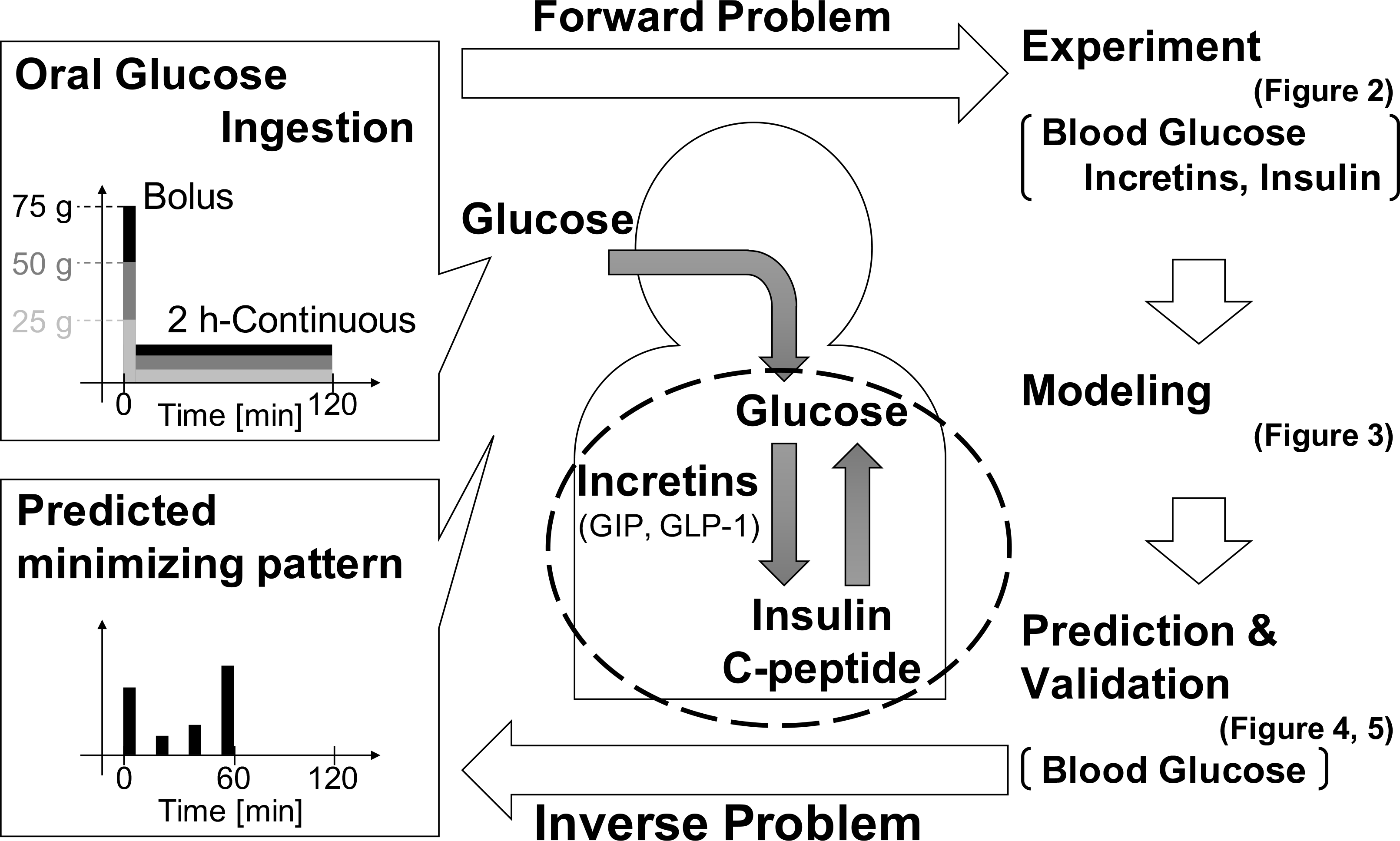
Study diagram. Three subjects orally ingested glucose with 3 doses 75 g, 50 g and 25 g in 2 durations of bolus and 2 h-continuous ingestion. Time course data of blood glucose level, insulin level, C-peptide level, GIP level, and GLP-1 levels were obtained (Figure 2). We constructed models of the dynamics of these blood hormones and glucose for each subject as a forward problem (Figure 3). Using the models, we predicted the minimization pattern, the glucose ingestion pattern minimizing the peak value of blood glucose level for the ingestion of 50 g glucose within 60 min as an inverse problem, and validated the pattern experimentally (Figure 4). To explore key parameters of the minimization pattern, we performed coarse-grain analysis (Figure 5).

With any ingestion pattern, the temporal pattern of each molecule exhibited a transient increase that returned to baseline within 4 hours (Figure 2). For bolus ingestion, the blood glucose and other blood hormones reached similar peak values for each dose of ingested glucose (Figure 2A, C, E, G, I). For the 2 h-continuous ingestion, blood glucose and other blood hormones showed increasing peak values with increasing doses of ingested glucose (Figure 2B, D, F, H, J). A consistent difference between bolus and continuous ingestion was that in the bolus ingestion case, with increasing doses of glucose, the time when blood glucose and hormones began to decrease and time to return to baseline become more delayed. In contrast, for 2 h-continuous ingestion, the time when blood glucose and other hormones began to decrease, and the time when all returned to the basal level were similar regardless of dose of ingested glucose. Subjects #2 and #3 showed similar responses to subject #1 by bolus and 2 h-continuous ingestion, except for GLP-1 (Figure S1). GLP-1 for only 75 g bolus ingestion for subject #1 showed a high transient peak, but that for subjects #2 and #3 did not.

**Figure 2.**
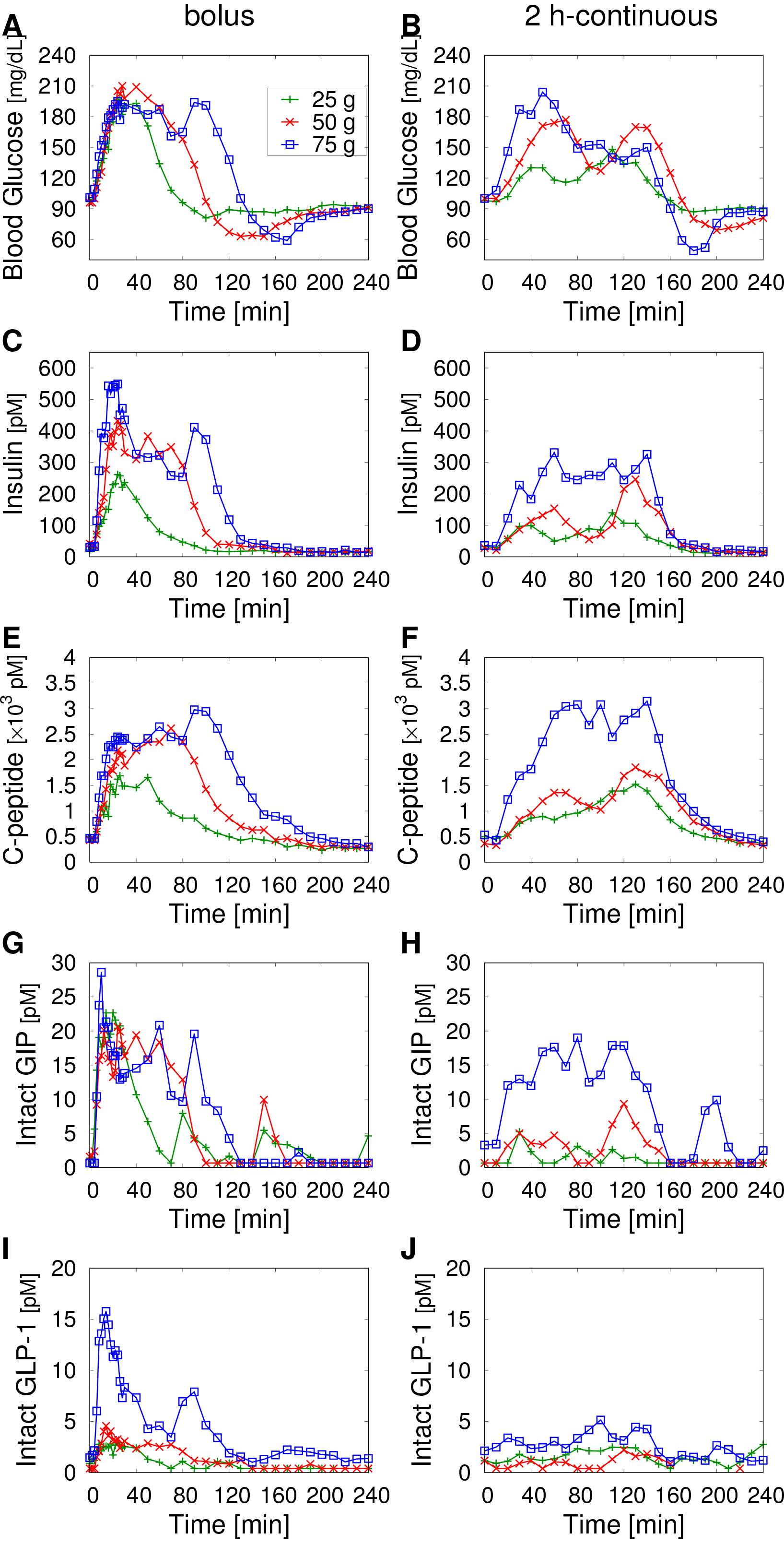
Time course data of blood glucose level and blood hormones in subject #1 by glucose ingestion. (A, B) Blood glucose. (C, D) insulin. (E, F) C peptide. (G, H) intact GIP. (I, J) intact GLP-1. (A, C, E, G, I) Bolus ingestion. (B, D, F, H, J) 2 h-continuous ingestion. The doses are indicated in panel A.

### Mathematical Model of Blood Glucose Control

As a solution to the forward problem, we constructed a mathematical model of blood glucose control that fits time course data of blood glucose and hormones. We constructed a mathematical model from ordinary differential equations (Figure 3A, Table 1, Table S1, see STAR Methods B.1). Because of possible alternative mechanisms of actions of GIP and GLP-1 on insulin secretion (Brubaker et al., 2007; Dalla Man et al., 2016; Kabul et al., 2015; Kim et al., 2014; Pedersen et al., 2011), we constructed multiple alternative models in which the GIP or GLP-1 or both have independent actions or cooperative actions with blood glucose to promote insulin secretion (Figure 3A, Table 1, Table S1, see STAR Methods B.1). We estimated parameters of each model for each subject separately to fit time course data of blood glucose and hormones. We selected the best model of blood glucose control for each subject by Akaike Information Criterion (AIC) ( see STAR Methods B.2). The selected models were the same for subjects #1 and #3, but different from the model for subject #2 (Table S2, S3). In the models of subject #1 and #3, cooperative action by blood glucose and GIP was selected, indicating that insulin secretion did not depend on GLP-1. In the model of subject #2, the independent action of GIP and Cooperative action by blood glucose and GLP-1 were selected. In each subject model, time course data of each blood glucose and hormones were approximately reproduced (Figure 3B, Figure S2, S3, Table S4).

**Table 1.**
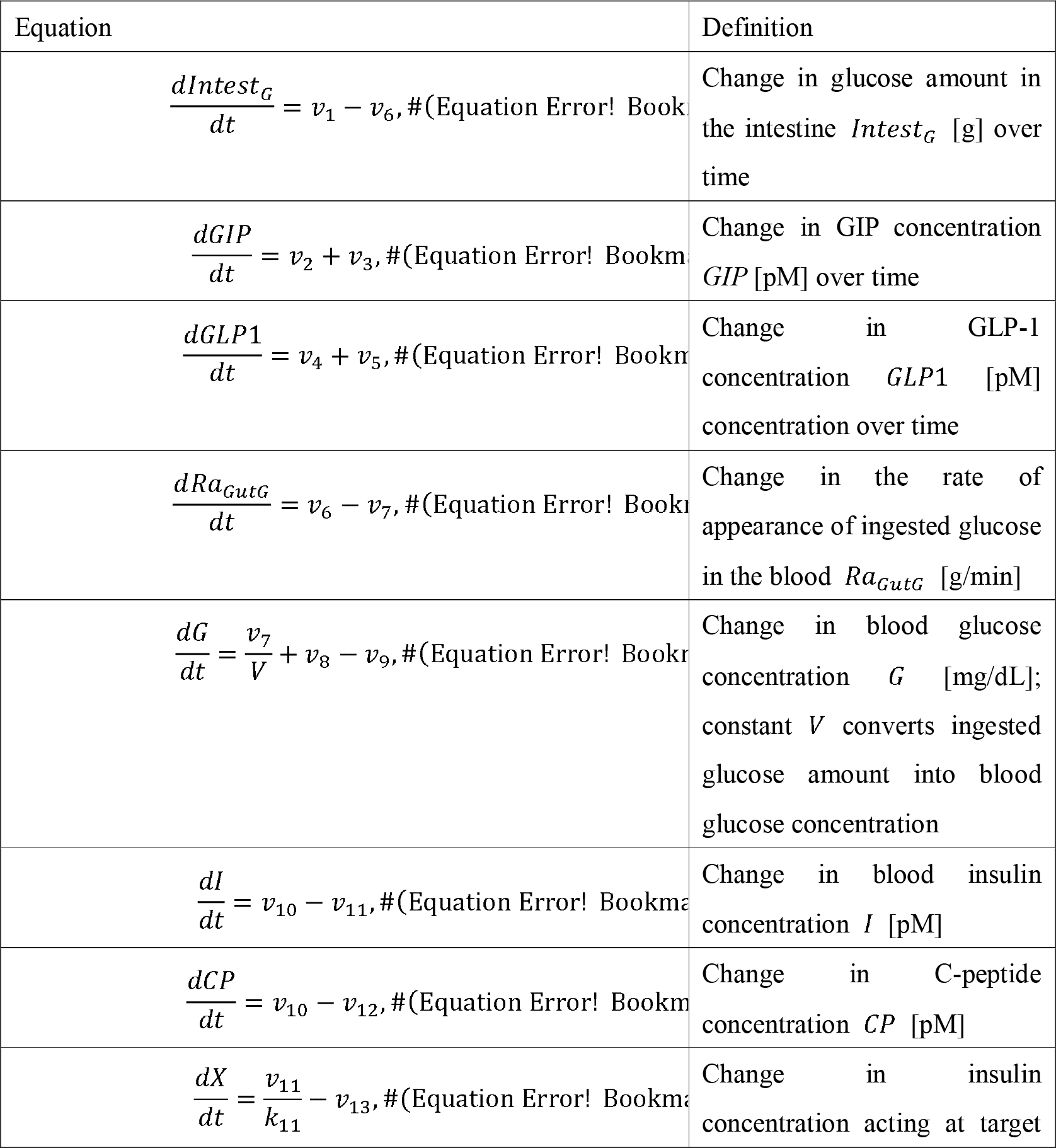
Equations in the ordinary differential equation models. The estimated parameters are the 18 parameters of *k*_2_,*k*_2_, *k*_4_, *k*_5_, *k*_6_, *k*_7_, *k*_8_, *k*_10_, *K*_2_, *K*_4_, *K*_6_, *K*_8_, *K*_11_, *V a,b,c*, and *d*. Simulations started from the models with initial concentrations of *GIP*(0),*GLP*1(0),*G*(0),*I*(0),*CP*, and *X*(0). Using the variables, and assuming *Duod*_*G*_, *Ra*_*GutG*_, *GIP*, *GIP*1, *G*, *I*, *X*, and *CP* are at steady state before ingestion, other initial conditions and parameters, *Ingest*_*G*_(0), *Ra*_*GutG*_(0), *k*_9_, *k*_11_, *k*_12_, *k*_13_, *GIP*_*B*_ and, *GIP*1_*B*_were determined by estimated parameters and initial values (Equations 23–30).

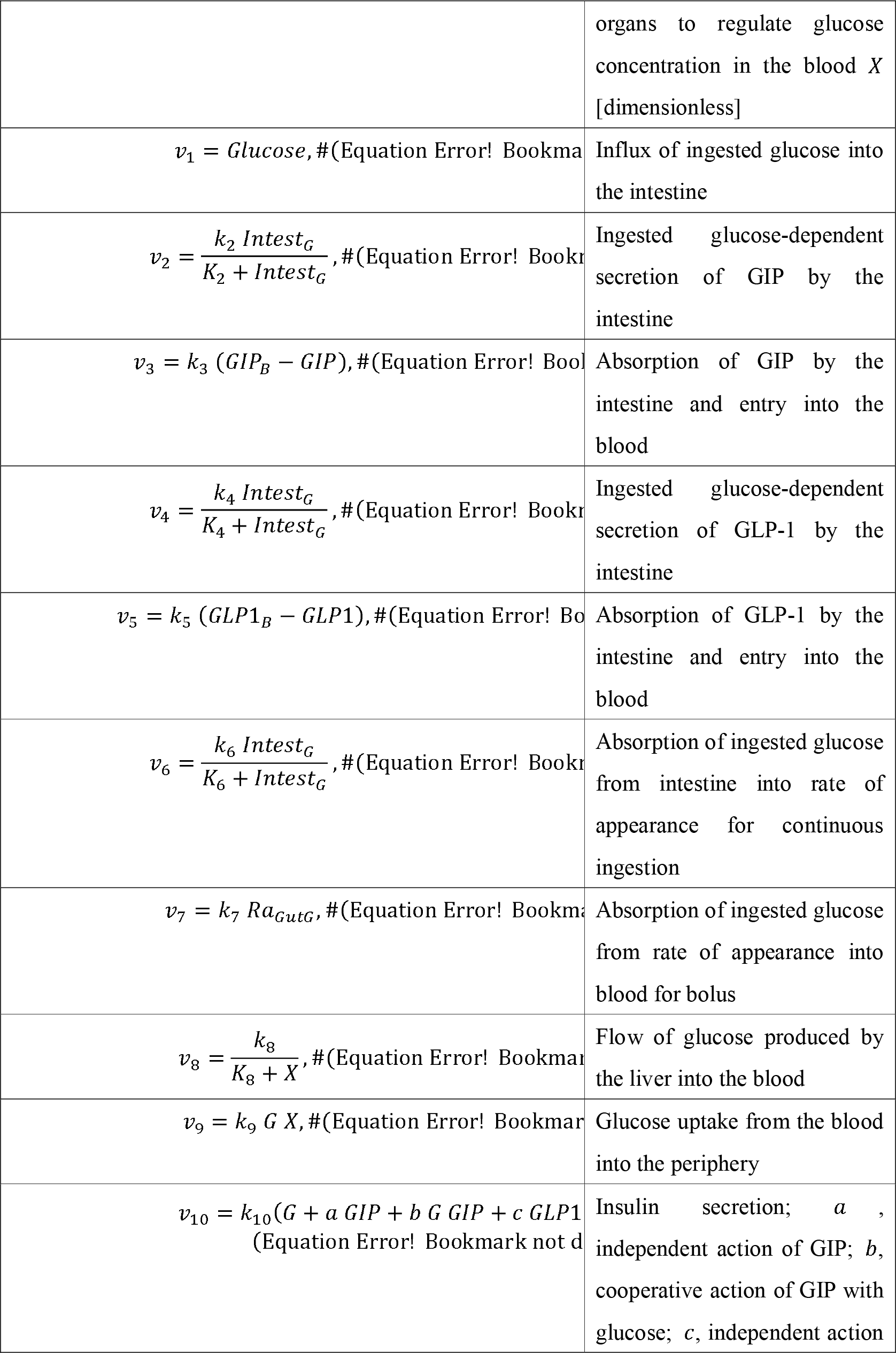

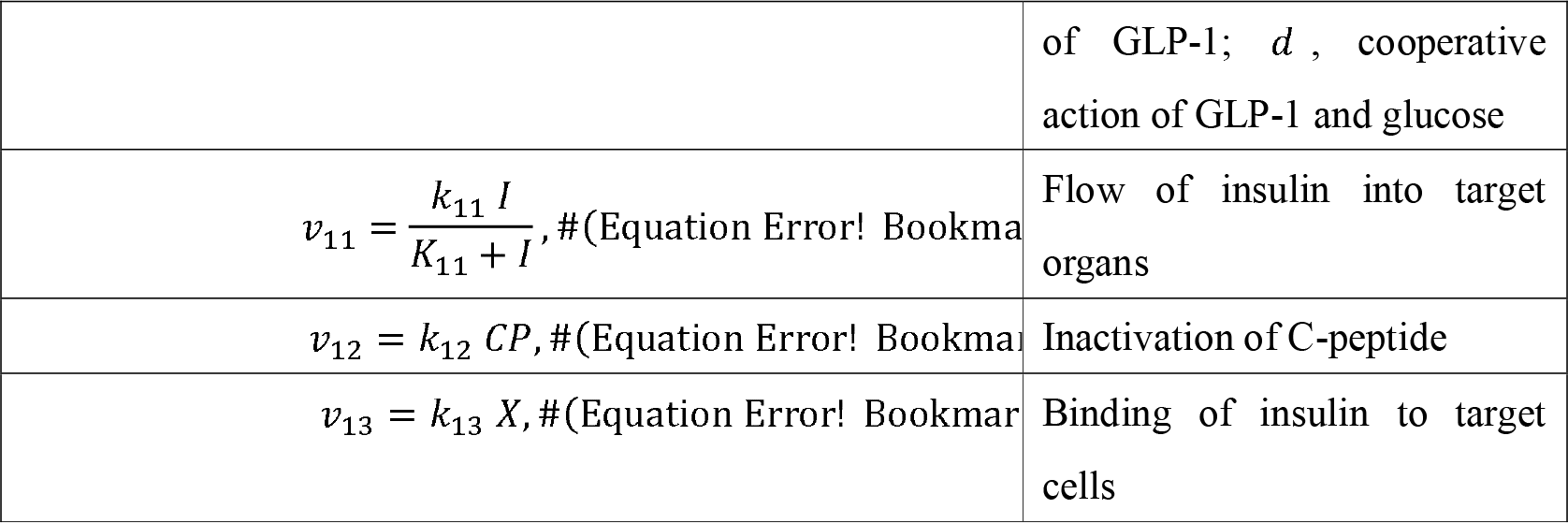

**Figure 3.**
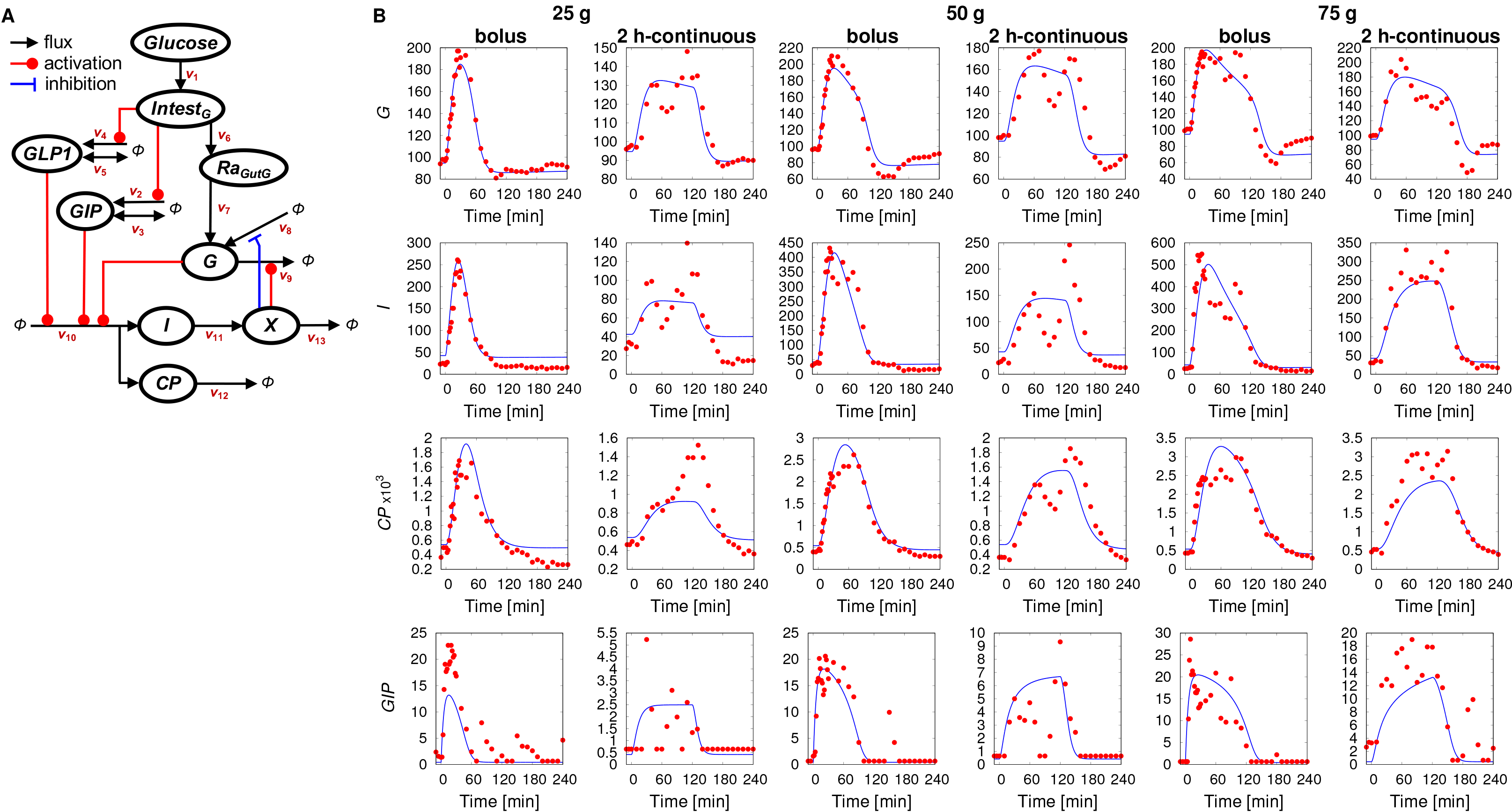
The blood glucose control model. (A) Model diagram. The letters in the circle indicate the variables of the model, the arrows indicate the flow of molecules, the red lines indicate activation, and the blue line indicates suppression (see STAR Methods B.1). The best fitting models for subjects #1 and #3 lack the GLP1 components. (B) Temporal patterns of hormones. The blue lines indicate the temporal patterns of simulations, and the red circles indicate the time course data of experiments. The dose and ingestion pattern are indicated at the top.

### Optimization and Validation of Glucose Ingestion Pattern that Minimizing Peak Value of Blood Glucose Level

Using mathematical, we tackled the inverse problem of predicting an optimal input pattern that optimally controls the output pattern. Here, input and output patterns are, specifically, time courses of oral glucose ingestion and blood glucose level, respectively. The optimality of the output pattern is defined as an objective function that is a function of the output pattern, typically the peak value of blood glucose level. First, we optimized the glucose ingestion pattern for each subject that minimized the typical objective function. Hereafter, we designate the optimized patterns minimizing objective function as the minimization pattern. We searched the solution under the following restrictions; total 50 g of glucose should be ingested within 60 min, glucose is ingested every 5 min, at least 1 g is ingested at 0 min and the remaining 49 g of glucose is distributed between 0 and 60 min. Because the combination of glucose ingestion patterns is enormous (62!/(49!12!)), we obtained an optimal ingestion pattern using an evolutionary programing-based optimization algorithm (see SART Methods B.3) (Back and Schwefel, 1993). The minimization patterns for the three subjects were designed with the above-explained method and shown in Fig. 4A.

**Figure 4.**
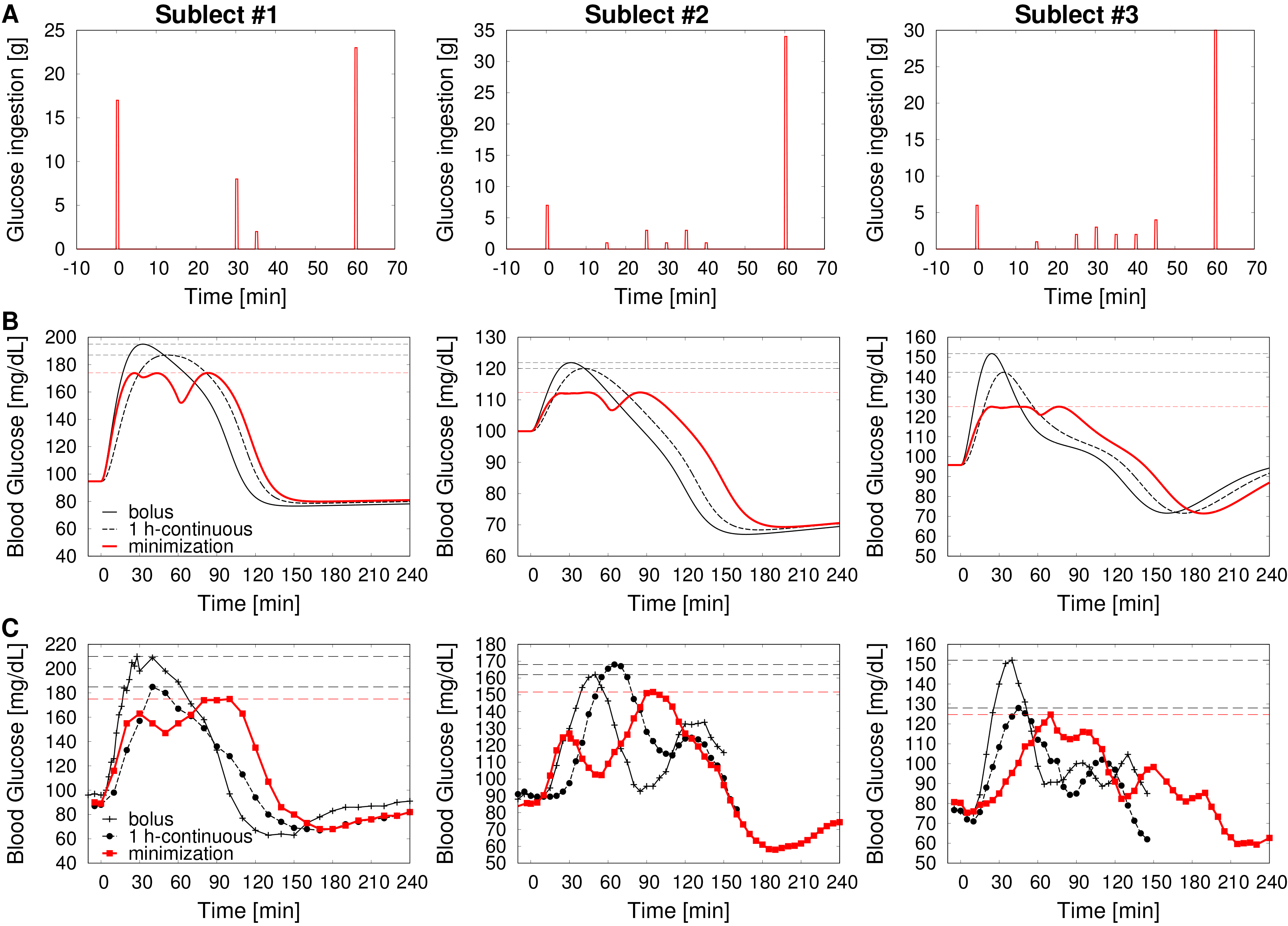
Optimal Patterns minimizing the peak value of blood glucose level. (A) Minimization patterns for glucose ingestion that minimize the peak value of blood glucose level in subject #1, #2, and #3. (B) Temporal patterns of blood glucose simulated from ingestion of glucose according to the minimization pattern (red line), bolus ingestion (black solid line), or h-continuous ingestion (black broken line). The peak values achieved for each ingestion pattern are marked with dashed horizontal lines. (C) Time course data of blood glucose level by the ingestion of the minimization pattern (red line and points and square symbols), the bolus ingestion (black solid line and open circles), and 1 h-continuous (black broken line and x symbols). The peak values achieved for each ingestion pattern are marked with dashed horizontal lines.

The optimized minimization pattern of the subject #1 appeared to be an intermittent pattern with 30-min intervals with most glucose ingested at 0 min (17 g) and 60 min (23 g), and smaller amounts ingested at 30 min (8 g) and 35 min (2 g) (Figure 4A, Table S5). This pattern was different from bolus and 1 h-continuous ingestions. The predicted blood glucose achieved with the minimization pattern showed a bimodal temporal pattern with peaks from ~25 min to 50 min and at ~80 min (Figure 4B, red line).

The optimized minimization patterns of subjects #2 and #3 appeared to be intermittent patterns similar to the pattern of the subject #1 (Figure 4A; Table S5). Compared with subject #1, for subjects #2 and #3, the optimized pattern of ingestion had some notable differences: Ingestion amount of glucose at 0 min was less, the number of time points at ~30-min the intermittent period during which glucose was ingested was larger, and the ingestion amount of glucose at 60 min was larger. The predicted blood glucose level achieved with the minimization pattern for subjects #2 and #3 showed a similar bimodal pattern to that for subject #1 (Figure 4B, red line).

We also compared the simulated blood glucose levels produced with the minimization pattern with those simulated for bolus or 1 h-continuous ingestion of 50 g of glucose. The predicted minimization pattern produced a lower peak value of blood glucose level than either simulations of bolus or 1 h-continuous ingestion using the subject-specific models (Figure 4B).

We validated the predicted blood glucose levels produced with the minimization patterns for each subject. Each subject ingested glucose according to their specific optimized minimization pattern (Table S5), and blood glucose levels were measured. (Figure 4C, red line). The peak value of blood glucose level produced by ingestion according to the minimization pattern in each subject was less than those produced by bolus and 1 h-continuous ingestion (Figure 4C, Table S6). All subjects exhibited bimodal temporal patterns of blood glucose level. These experimental results are consistent with the predictions except that first peak in blood glucose level at ~30 min was lower than the second peak at ~80 min for subjects #1 and #2 and the peak in blood glucose was delayed from the prediction for subject #3.

To examine how the key parts of the ingestion pattern that resulted in the pattern that minimized the peak value of blood glucose level, we simplified the ingestion pattern into a coarse-grained ingestion pattern (Figure 5). We generated a minimization pattern with 5 parameters (Figure 5A, see STAR Methods B.4), and examined the effect of parameters on the peak value of blood glucose level.

**Figure 5.**
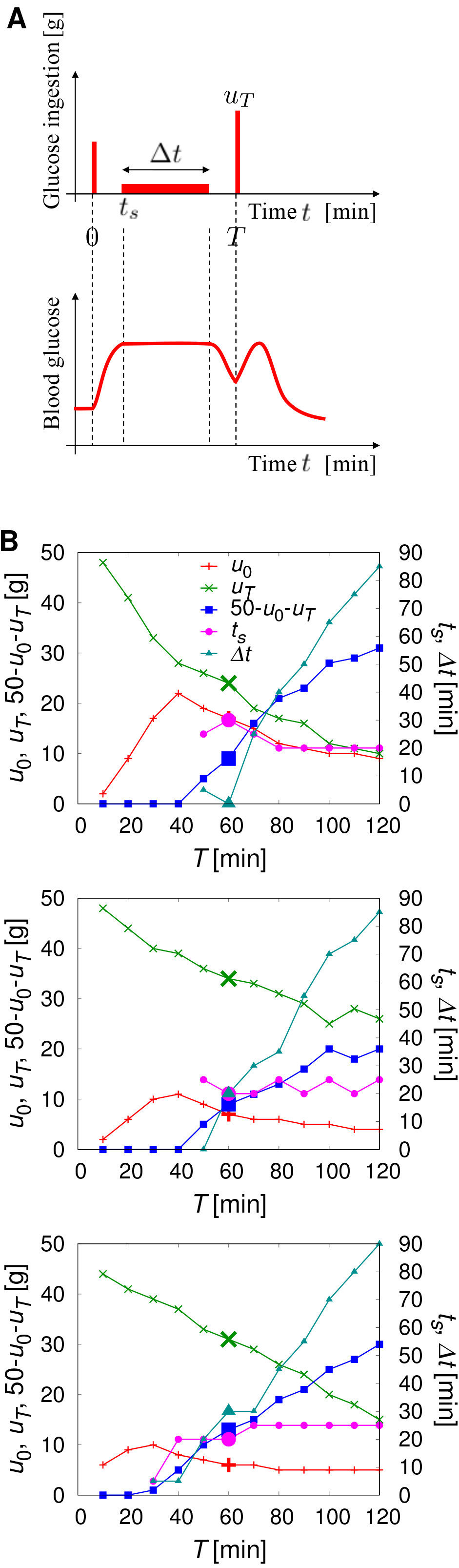
Coarse-grain analysis of minimization pattern. (A) Coarse-grained minimization pattern characterized by 5 parameters, the dose of ingested glucose at 0 min *u*_0_, the start time of the intermediate period *t*_*s*_, the duration of intermediate period *Δt*, the dose of ingested of glucose at the end time *u*_*T*_, and the end time of the ingestion *T* (B) *T*-dependency of *u*_0_, *u*_60_, *t*_s_, and *Δt* that realize the minimum value of peak value of blood glucose level for each subject.

We coarse-grained the minimization pattern into three periods; start time (0 min) of the first bolus ingestion, continuous-like intermediate period, and the end time of the ingestion (Figure 5A, upper panel). The coarse-grained pattern was characterized by 5 parameters; the dose of ingested glucose at 0 min *u*_0_, the start time of the intermediate period *t*_*s*_, the duration of intermediate period *Δt* the dose of ingested of glucose at the end time *u*_*T*_, and the end time of the ingestion *T* (Figure 5A, see STAR Methods B.4). We changed the parameters, and examined the effect of each parameter on the peak value of blood glucose level (Figure 5B).

To determine the parameter sets of the coarse-grained minimization pattern, we fixed *T* = 60 [min], the same duration as Figure 4A, and identified the parameter set that minimized the peak value of blood glucose level of subject #1 (Figure 5B, larger symbols). The parameter set *u*_0_ = 17 [g], *u*_60_ = 24 [g], *t*_*s*_ = 30 [min], *Δt* = 0 [min] produced a minimum (174.07 mg/dL) value for the peak value of blood glucose level, and this value is equivalent to that (173.95 mg/dL) achieved with the minimization pattern (Table 2). The coarse-grained minimization pattern was almost the same as that of the minimization pattern of subject #1 obtained in Figure 4. This result indicates that the coarse-grained minimization pattern captures essential characteristics of the minimization pattern, such as the peak value of blood glucose level and the temporal pattern.

**Table 2.** Properties of the models. The ODE model has 16 parameters. The roles of the incretins in each subject’s best fitting full model are shown, along with the predicted peak of blood glucose concentrations achieved with the minimization. The values of the parameters of the coarse-grained models that produced the peak value of blood glucose level for each subject are indicated.

|  | ODE model | Course-grained minimization pattern |
| --- | --- | --- |
| Subject #1 | Cooperative effect of GIP with glucose<br>Minimization pattern: peak value of blood glucose level = 173.95 mg/dL | $u_0 = 17$ [g]<br>$u_{60} = 24$ [g]<br>$t_s = 30$ [min]<br>$\Delta t = 0$ [min]<br>Peak value of blood glucose level = 174.07 mg/dL |
| Subject #2 | Independent effect of GIP<br>Cooperative effect of GLP-1 with glucose<br>Minimization pattern: peak value of blood glucose level = 112.36 mg/dL | $u_0 = 7$ [g]<br>$u_{60} = 34$ [g]<br>$t_s = 20$ [min]<br>$\Delta t = 20$ [min]<br>Peak value of blood glucose level = 112.43 mg/dL |
| Subject #3 | Cooperative effect of GIP with glucose<br>Minimization pattern: peak value of blood glucose level = 125.14 mg/dL | $u_0 = 6$ [g]<br>$u_{60} = 31$ [g]<br>$t_s = 20$ [min]<br>$\Delta t = 30$ [min]<br>Peak value of blood glucose level = 125.75 mg/dL |

We examined the dependency of the parameters on the end time of ingestion *T* for minimizing the peak value of blood glucose level (Figure 5B, top). When *T* ≤ 40 [min], *u*_0_ + *u*_*T*_ = 50 [g], and, as *T* increases *u*_0_ increases (Figure 5B, red line), and *u*_*T*_ decreases (Figure 5B, green line). Thus, when the total ingestion time is within 40 min, (i) there is no intermediate period, and (ii) as the total ingestion time increases, the dose of ingested glucose at 0 min increases and the dose of ingested glucose at the end time decreases. When *T* > 40 [min], both *u*_0_ and *u*_*T*_ decrease as *T* increases, and the dose of ingested glucose at the intermediate period 50 − *u*_0_ − *u*_*T*_ increases almost linearly (Figure 5B, blue line). Thus, as the total ingestion time is longer than 40 minutes, (i) the dose of ingested glucose both at 0 min and at the end time decrease and (ii) the dose ingested during the intermediate period increases. Also, when *T* > 40 [min], *t*_*s*_ is almost constant between 20 [g] and 30 [g] (Figure 5B, magenta line). Δ*t* is nearly 0 [min] when *T* ≤ 60 [min], and increases almost linearly when *T* > 60 [min] (Figure 5B, cyan line). Together these simulations indicated that, when the total ingestion time is shorter than 60 min, the duration of the intermediate period becomes so short that ingestion becomes bolus-like, and the intermediate period becomes longer as the total ingestion time is longer than 60 min. For subjects #2 and #3, the dependency of the parameters on the end time of ingestion *T* was also qualitatively the same (Figure 5B, middle and bottom).

Changing the duration of intermediate period *Δt* caused the greatest differences among the subjects (Figure S4). When *T* = 60 [min], the duration of the intermediate period of subject # 1 is shorter than those of subject #2 and #3, which was also shown in non-coarse-grained minimization patterns for the subjects (see Figure 4). For both subjects #2 and #3, as the total ingestion time becomes shorter than 60 min, the duration of the intermediate period becomes shorter and ingestion becomes bolus-like as for subject #1 (Figure 5B, Figure S4). Thus, the length of the total ingestion time is predicted to control the duration of the intermediate period among the subjects with all subjects exhibiting a point at which a bolus-like ingestion occurs during the intermediate period.

The quantitative difference of duration of intermediate period among subjects may relate to differences in glucose tolerance among subjects. Glucose tolerance is determined by the balance between insulin secretion, sensitivity, and clearance; as glucose tolerance decreases, insulin secretion, sensitivity, and clearance also decrease (Antuna-Puente et al., 2011; Ohashi et al., 2018, 2015; Polidori et al., 2016; Schofield and Sutherland, 2012). Therefore, we examined the effect of the parameters for the reaction rates for insulin secretion, sensitivity and clearance on duration of the intermediate period (Figure S5). The models for each subject showed that the duration of the intermediate period becomes longer as the insulin secretion or sensitivity increase and becomes shorter as the insulin clearance increases. Thus, the duration of the intermittent period does not correspond to these insulin-related parameters controlling glucose tolerance.

## DISCUSSION

### Prediction and Validation of Glucose Ingestion Patterns that Minimize the Peak Value of Blood Glucose Level

In this study, as a forward problem, we constructed a mathematical model of the change in blood glucose from time course data of blood glucose and hormones in blood during and following oral glucose ingestion with various doses and durations in human subjects. Using this model, as an inverse problem, we optimized glucose ingestion patterns that minimize the peak value of blood glucose level and validated these patterns with the human subjects by experiments. The minimization pattern was an intermittent pattern different from both the bolus ingestion and the continuous ingestion. This intermittent ingestion pattern was intuitively not obvious. However, we discovered the pattern using this approach of both constructing a mathematical model as a forward problem and optimizing input pattern from the model as an inverse problem. Although the best fitting model for each subject had important differences in the roles of the blood hormones, the intermittent pattern as an optimal ingestion pattern to minimize peak value of blood glucose level was common to all three subjects, suggesting that the minimization pattern is robust to these differences in the model. Although we determined that the duration of the intermittent period was a key parameter controlling the minimization pattern output, we did not determine a molecular mechanism for how the intermittent pattern minimizes the peak value of blood glucose level. This question will be analyzed in the future.

Methodologically, construction of a mathematical model based on the experimental data as a forward problem is well-known. However, the inverse problem of optimizing an input pattern to achieve a specified output pattern is challenging (Murakami et al., 2017). Our success in identifying optimal input patterns through analysis of both the forward problem and inverse problem suggests that this approach is valid for biological systems. An obvious potential application is designing optimal ingestion patterns for various nutrients or combinations of nutrients such that the ingestion pattern that minimizes the peak value of blood glucose level can be logically designed. Such logical design of optimal food ingestion pattern will contribute the human metabolic care and the prevention of the type 2 diabetes. Here, the objective function for which we predict the input pattern is the peak value of the blood glucose level. By changing the objective function, this approach can evaluate other biological outputs and predict the input pattern that optimizes molecular concentrations or other measurable factors.

### Identification of Individualized Models of the Control of Blood Glucose Level

Our ordinary differential equation models include the roles of incretins in insulin secretion. By determining the best fitting model for each subject, we observed differences between subjects in the roles of incretins in regulating blood glucose level. None of the subjects had models that included an independent effect of GLP-1 on insulin secretion. Two of the three subjects had no role for GLP-1 (independent or cooperative with glucose) in their optimal models. In previous mathematical models using Caucasians data, only GLP-1, but not GIP, were incorporated (Dalla Man et al., 2016; Pedersen et al., 2011). It has been reported that secretion of intact GLP-1 in Japanese is very small, although that of the total GLP-1 in Japanese is almost the same as that in Caucasians (Seino et al., 2010). All subjects in this study are Japanese, and, the intact GIP level was higher than the intact GLP-1 level for all of them (Figure 2). Intact GLP-1 and intact GIP have a similar EC_50_ for their receptors: The EC_50_ of intact GIP is 8 nM (Gespach et al., 1984), and the EC50 of intact GLP-1 is 2.6 nM (Adelhorst et al., 1994). Considering the higher level of intact GIP than intact GLP-1 in the blood and their similar sensitivities, it is reasonable that intact GIP rather than intact GLP-1 was the incretin with the most effect on insulin secretion in the best fitting model.

Many mathematical models use average values of blood glucose from many subjects of all subjects. Some models that use data from individual subjects used data with only a single dose of glucose (Dalla Man et al., 2013; De Gaetano et al., 2013; Ohashi et al., 2018, 2015). Here, we used data from individual subjects using 3 different doses and 2 different durations of glucose ingestion. We constructed a mathematical model using a single dose of glucose (75 g, like that of the OGTT) in subject #1 and compared this OGTT 327 model with the model that we constructed from the data for the 3 different doses and 2 different durations of glucose ingestion (Figure S6). The model that used the multiple dose and ingestion durations had a better fit to the blood glucose level achieved by ingestion of glucose according to the minimization pattern (lower *RSS* value) than did the model using 75 g OGTT alone. Thus, the single dose OGTT appears insufficient to reflect the dynamics of the blood glucose level in sufficient detail for mathematical modeling, and models should be constructed from data on multiple doses and durations of glucose ingestion to be useful in predicting the minimization pattern.

This finding that more training data provides more accurate predictive power is expected. However, the number of conditions for training data sets is limited in humans, because these types of studies take a long period of time and require several hours and fasting by the participants for each experimental condition. Here, we set an interval of 1 to 2 months for each experiment, thus collecting the data required a minimum of six months, and, in reality, more than a year. During such a long period, having many subjects for blood sampling following oral glucose ingestion every month over a year is difficult. Changes in the state of a subject can change during the months of the experiment, which can affect the model and reduce predictive power. Thus, in human subject tests, there is a trade-off relationship between the number of training data sets and the prediction accuracy.

A limitation of the study is that the model is limited. There are mechanisms, such as glucagon, autonomic nerves, and free fatty acids, that did not incorporate into the model. Glucagon is a counter-acting hormone to insulin in regulation of blood glucose. Glucagon increases blood glucose level by facilitating glycogenolysis (Alberti and Zimmet, 1998; Jiang and Zhang, 2003). Autonomic nerves not only regulates secretion of insulin and glucagon (Thorens, 2011), but also affects hepatic glucose production and uptake (Kimura et al., 2016; Ruud et al., 2017). Free fatty acids weakens the effect of insulin on hepatic glucose production and peripheral glucose uptake (Okuno et al., 1998; Yamauchi et al., 2001). Although glucagon and free fatty acid are not explicitly incorporated in our model, blood glucose levels and other hormone concentrations are well reproduced, which may suggest that the effects of other molecules such as glucagon and free fatty acid, are implicitly incorporated by some parameters in the model. Incorporating glucagon and free fatty acid explicitly into the model is a future goal.

In conclusion, the key points of this study are three. The first point is the experimental design. We performed six different conditions of oral glucose ingestion (3 doses and 2 durations) for each subject and obtained detailed time course data, which makes the model predictable. The second point is the demonstration of ability to logically design blood glucose control. We predicted and validated the oral glucose ingestion pattern that minimized the peak value of blood glucose level. The third point is the methodology. We solved a forward problem by constructing the mathematical model of output with the given input patterns, and in turn, solved an inverse problem by logical designing the input pattern to control the output pattern. We expect that this approach with a forward problem and an inverse problem that are solved using the mathematical model can be widely applied to design optimal dietary ingestion pattern relevant to human health.

## STAR⋆Methods

Detailed methods are provided in the online version of this paper and include the following:

- METHOD DETAILS

- A. Experiment
- B. Model and Analysis

## SUPPLEMENTAL INFORMATION

Supplemental Information includes eight figures and seven tables.

## AUTHOR CONTRIBUTIONS

Conceptualization and Methodology, M.F., Y.M., S.O. and S.K.; Experiment, Y.K., Y.S., and S.F.; Modeling and Simulation, M.F., and Y.M.; Analysis, M.F., Y.M., M.K., S.U., H.K., H.I., K.K., S.O., S.I, S.K.; Writing, M.F., Y.M., Y.K., and S.K.; Supervision and Funding, S.I. and S.K.

## ACKNOWLEDGEMENT

We thank fellow laboratory members for critical reading of the manuscript and for technical assistance with the experiment and analysis. We thank Rika Sumita, Mina Shiguma, and Naomi Isene for technical assistance. We thank Nancy R. Gough (BioSerendipity, LLC) for editing the manuscript. The computations for this work were performed in part on the NIG supercomputer system at ROIS National Institute of Genetics. This work was supported by the Creation of Fundamental Technologies for Understanding and Control of Biosystem Dynamics (JPMJCR12W3), CREST, of the Japan Science and Technology Agency (JST) and by the Japan Society for the Promotion of Science (JSPS) KAKENHI Grant Number (17H06300, 17H6299, 18H03979). M.F. receives funding from a Grant-in-Aid for Challenging Exploratory Research (16K12508). Y.K. receives funding from a Grant-in-Aid for Young Scientists (18K16578). S.U. receives funding from a Grant-in-Aid for Scientific Research on Innovative Areas (16H01551) and a Grant-in-Aid for Scientific Research on Innovative Areas (18H04801). H.K. receives funding from a Grant-in-Aid for Scientific Research on Innovative Areas (16H06577). K.K. receives funding from a Grant-in-Aid for Scientific Research (B) (15KT0021) and (C) (15K00246), and a Grant-in-Aid for Scientific Research on Innovative Areas (16H01554). The funders had no role in study design, data collection and analysis, decision to publish, or preparation of the manuscript.

## STAR⋆Methods

### KEY RESOURCE TABLE

| REAGENT or RESOURCE | SOURCE | IDENTIFIER |
| --- | --- | --- |
| Software and Algorithms |  |  |
| Ordinary Differential Equation Simulations and Analysis - MATLAB | Mathworks | RRID:SCR_001622 |

### METHOD DETAILS

#### A. Experiment

##### A.1. Subjects

The subjects’ profiles are as shown in Table S7. All subjects are healthy, and signed informed consent.

##### A.2. Blood Sampling Experiment

For oral glucose tolerance test, a glucose solution containing 25 g, 50 g or 75 g glucose was orally ingested after a 10-hour fasting, and blood samples were obtained at the times indicated in the figures from the cutaneous vein of the forearm. Blood samples were obtained from the cutaneous vein of the forearm. Blood collection on fasting was performed twice and then a glucose solution containing 25 g, 50 g or 75 g glucose was orally ingested. The ingestion method was rapid within a minute (bolus ingestion), and continuous over the course of 2 hours (2 h-continuous ingestion). For continuous ingestion, we connected the tube to noncontact microdispenser robot (Mr. MJ; MECT Corporation) (Sano et al., 2016) and glucose solution was ingested from tube. To equalize the volume of ingested glucose solution, glucose solution, TRELAN-G75 (AJINOMOTO), was diluted with distilled water into a total volume 225 ml. Each amount of glucose and delivery paradigm was tested with each subject in experiments separated by at least 1 month. Blood was rapidly centrifuged, plasma glucose and hormone concentrations expect for GIP were measured according to the methods with LSI Medience Co., Ltd. Plasma glucose was measured by enzymatic methods (IATRO LQ GLU). Plasma insulin and Serum C-peptide was measured by Chemiluminescent Immunoassay (Tholen et al., 2004; Tietz, 1990). Plasma intact GLP-1 and Plasma intact GIP were measured by ELISA kits (#EGLP-35K, Merck, Billerica, MA or #27201, Immuno-Biological Laboratories, Gunma, Japan, respectively) (Miyawaki et al., 2002; Tijssen, 1985). For simplicity, we refer to plasma glucose, plasma insulin, serum C-peptide, plasma intact GIP, and plasma intact GLP-1 as blood glucose, insulin, C-peptide, GIP, and GLP-1, respectively.

##### A.3. Validation Experiment

For the validation experiment of the minimization pattern, we employed the same method as described in A.2 for the subject #1, and a Freestyle Libre continuous glucose monitoring system (FGM; Abbott Diabetes Care) for subjects #2 and #3. FGM reduces the invasive burden on the subjects because the subjects wear a sensor rather than requiring an indwelling needle for blood glucose monitoring. We performed the experiment after the subject had worn the sensors for at least two days. Each subject wore three sensors, and bolus ingestion, continuous ingestion for 1 hour, ingestion of minimization pattern were carried out using the same sensors within two weeks. The results of the three sensors were averaged for each paradigm. Because FGM measures glucose level of the interstitial fluid rather than glucose level in the blood, the measured value reflects a delay of about 5 to 20 minutes (Figure S3) compared with the values obtained by blood collection.

##### A.4. Ethics Committee Certification

We complied with Japan’s Ethical Guidelines for Epidemiological Research, and the study as approved by the ethics committees of the Life-Science Committee of the University of Tokyo (16-265). Subjects were recruited by the related law.

#### B. Model and Analysis

##### B.1. Model Structure and Parameter Structure

For each subject, we estimated parameters that reproduce the time course data of blood glucose, insulin, C-peptide, intact GIP, and intact GLP-1 of six glucose ingestion patterns, combinations of three doses (25 g, 50 g, and 75 g) and given by bolus and 2 h-continuous ingestion, using the following model (Equations 1–21, Table 1).

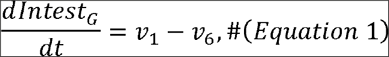

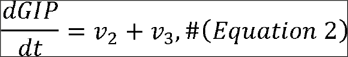

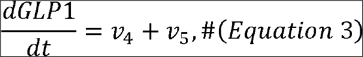

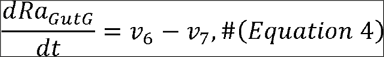

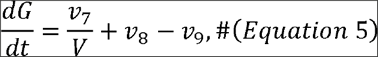

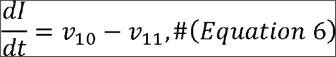

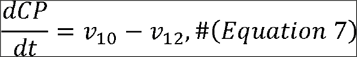

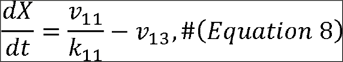

Equations 1-8 indicate differential equations reproducing time developments of glucose amount in the intestine *Intest*_*G*_ [g], GIP level *GIP* [pM], GLP-1 level *GLP*1 [pM], rate of appearance of ingested glucose amount into the blood *Ra*_*GutG*_ [g/min], blood glucose level *G* [mg/dL], insulin level *I* [pM], C-peptide *CP* [pM], and the insulin level acting on the regulation of glucose *X* (denoted as effective insulin concentration at target organs hereafter). Each variable is controlled by fluxes *ν*_*i*_ {*i* = 1,…,13}. However, in Equation 5, *ν*_7_ was divided by the constant *V* to convert the ingested glucose amount into the blood glucose level. Also in Equation 8, *ν*_11_ was divided by *k*_11_ to render *X* dimensionless. Rendering *X* dimensionless enables the elimination of redundant parameters, and improves the accuracy of parameter estimation. The fluxes *ν*_*i*_ are given by

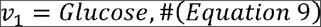

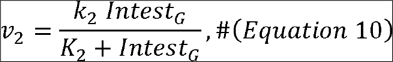

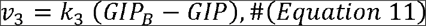

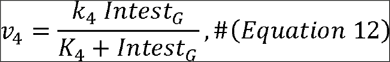

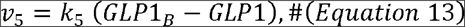

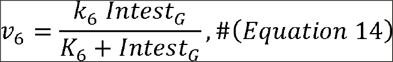

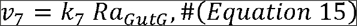

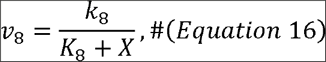

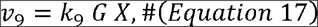

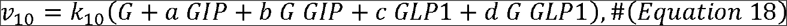

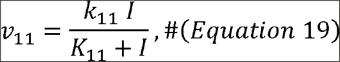

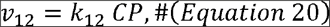

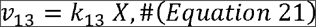

(Table 1, Figure 3).

*ν*_1_ indicates the influx of ingested glucose into the intestine, given by dose of glucose ingestion divided by the time duration of ingestion *Δt* otherwise 0 (Equation 22). For rapid ingestion, such as bolus ingestion, or for the ingestion of minimization pattern, *Δt* is assumed as 0.5 [min]. For example, in the case of 50 g bolus,

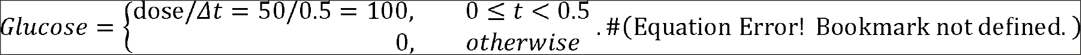

*ν*_2_ indicates the secretion of GIP depending on the glucose amount in the intestine (*Intest*_*G*_). *ν*_3_ indicates absorption of GIP by the intestine and entry into the blood, which is proportional to *GIP* subtracted by its basal *GIP*_*B*_. At steady state without glucose ingestion, *GIP* converges to *GIP*_*B*_. *ν*_4_ indicates the secretion of GLP-1 depending on the glucose amount in the intestine. *ν*_5_ indicates the absorption of GLP-1 proportional to *GLP*1 subtracted by its basal *GIP*1_*B*_. At the steady state without glucose ingestion, *GLP*1 converges to *GLP*1_*G*_. *ν*_6_ indicates the flow of glucose from the intestine to the rate of appearance (*Ra*_*Gutc*_). With bolus ingestion, this flow can be regarded as constant because of the large amount of glucose in the intestine (Brubaker et al., 2007). Therefore, we assumed that this flux is given by the Michaelis-Menten equation, which saturates when the glucose amount is large. *ν*_7_ indicates the flow of glucose from the rate of appearance into the blood, which is proportional to the rate of appearance of ingested glucose amount. *ν*_8_ indicates the flow of glucose production from the liver into the blood, given by an inhibitory Michaelis-Menten equation, which decreases as the amount of effective insulin *X* increases. *ν*_9_ indicates the glucose uptake from the blood to the periphery and is given by the product between blood glucose level *G* and effective insulin *X*.*ν*_10_ indicates the secretion of insulin. In this study, the actions of GIP and GLP-1 on insulin secretion were represented as independent actions of each incretin and as cooperative actions with blood glucose. By incorporating the parameters (*a,b,c,* and *d* in Equation 18), we could relate insulin secretion to cooperative or independent actions using AIC (Akaike Information Criteria) to select the model that best fit the data (Table S1, S2). *ν*_11_ indicates the flow of insulin *I* into target organs, such as liver and muscle, leading to effective insulin *X*. *ν*_12_ indicates inactivation of C-peptide *CP* and decreases in proportion to *CP* itself. *ν*_13_ indicates the binding of *X* to the cells in the target organs in proportion to *X* itself.

For the model, parameters were estimated for each subject. Here, the estimated parameters are the 18 parameters of *k*_2_,*k*_3_,*k*_4_,*k*_5_,*k*_6_,*k*_7_,*k*_8_,*k*_10_,*K*_2_,*K*_4_,*K*_6_,*K*_8_,*K*_11_,*V,a,b,c* and *d*; and six initial levels of *GIP*(0), *GLP*1(0), *G*(0),*I*(0),*CP* and *X*(0). Using the variables, and assuming *Duod*_*G*_, *Ra*_*GutG*_, *GIP*, *GIP*1, *G*, *I*, *X*, and *CP* are at steady state before ingestion, other initial conditions and parameters were determined by estimated parameters and initial values, given by

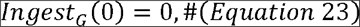

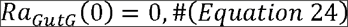

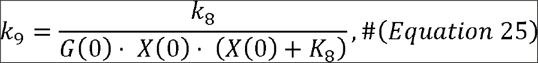

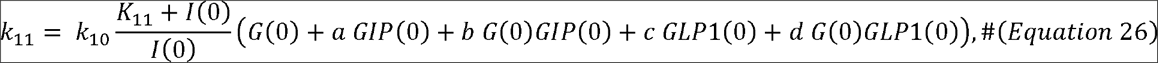

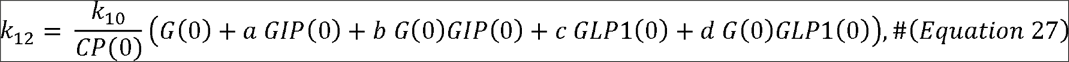

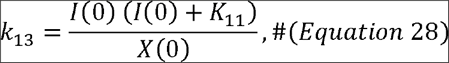

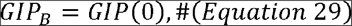

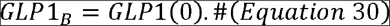

These parameters are different between subjects, but the same for each subject for each experimental paradigm (dose and duration and ingestion). This means that the state for each subject does not change during this study. For time development, we used CVODE in Matlab’s Systems biology toolbox.

We used the residual sum of squares as the objective function so that the residual between the experimental value and the simulation value is reduced, given by

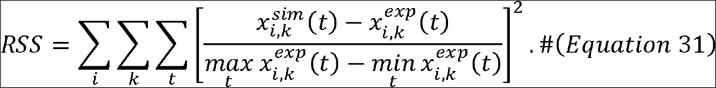

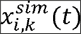 and 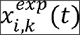 indicate the simulation values and the experimental values of molecular species *k* ∈ {*G,I,CP,GLP*1, *GIP*} at time *t* in the experiment *i* ∈ {25B,25C,50B,50C,75B,75C}, for which each experiment is denoted by the ingestion dose and the initial letters of the duration of ingestion, 25 g-bolus ingestion, 25B and for 75 g-2 h-continuous, 75C. To avoid the influences of the differences in the absolute quantities of the molecules, we normalized the difference between the simulation value and the experimental value by the difference between the maximum value and the minimum value of the experiment.

We performed parameter estimation for global optimal solution using Evolutionary programming (Bäck and Schwefel, 1993) for 40 trials with a parent number of 5000 and a generation number of 5000, then we obtained a local optimal solution using the simplex search method (Matlab fminsearch). We implemented all programs using Matlab 2015a and performed parameter estimation using 2.6 GHz CPU (Xeon E5 2670) at the National Institute of Genetics (NIG), Supercomputer System of Research Organization of Information and System (ROIS).

##### B.2. Model Selection

Using parameters of *a,b,c* and *d* in Equation 18, which indicate contributions to insulin secretion of incretins as independent actions of each incretin and cooperative actions with glucose, we considered the multiple models shown in Table S1.

We performed the parameter estimation of each of the above models using *RSS* of Equation 31 for each subject. Here, we assumed that each residual of the simulation value and the experiment value in Equation 31 follows a normal distribution. Among the models to be compared, the sum *N* of the numbers of data of each variable measured in the experiment is the same. Therefore, *AIC* (Akaike Information Criteria), which is a criterion of model selection can be calculated for each model, given by

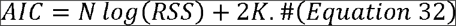

We employed a model that minimizes AIC for each subject as a model representing the dynamics of blood molecules in the subject. For the models not including GLP-1 of subjects #1 and #3 as mentioned below, we also cAICulated *AIC* for each model similar to those including GLP-1.

The selected models for each subject were distinct (Table 2, Table S1, S2). For subject #1, the best model had no influence of GLP-1 and both an independent action and cooperative action with glucose for GIP (Table S2, *c* = *d* = 0), indicating that the insulin secretion of subject #1 is independent of GLP-1. For subject #2, the best model had an independent action of GIP and a cooperative action of GLP-1 with glucose (Table S2, *b* = *c* = 0), indicating that the insulin secretion of subject #2 depends on both GIP and GLP-1. For subject #3, the best model had only the cooperative action of GIP with glucose (Table S2, *a* = *c* = *d* = 0), indicating that the insulin secretion of subject #3 is independent of GLP-1. In each subject model, time course data of each blood glucose and hormones were approximately reproduced (Figure 3B, Figure S2, S3).

In the selected models of subjects #1 and #3, insulin secretion did not depend on GLP-1, therefore, we performed parameter estimation and model selection using models that did not include GLP-1 by removing Equation 3. Insulin secretion using the best fitting of these models for both subjects #1 and #3 included the term of independent action of blood glucose and the cooperative term of blood glucose and GIP (Table S3, *a* = 0). We used these models for subjects #1 and #3.

##### B.3. Estimation of Minimization Patterns

We set the oral glucose *u*(*t*) as a function of time *t* [min] according to the following constraint condition. First, glucose was orally ingested at intervals of 5 min from 0 min to 60 min. Here, we defined *u*_*s*_ [g] as the dose of ingestion at the minute *s* [min] (*s* = 0,5,…, 60) and *u*_0:60_ as the temporal pattern of oral glucose ingestion, given by

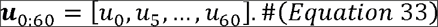

Also, we set the total dose of glucose ingestion at 50 g, *i.e.* Σ_s_*u*_*s*_ = 50 [g], each dose at *s* is the integer value with unit of 1 g, *i.e*. *u*_*s*_ ∈ 𝕫, *u*_*s*_ ≥ 0, and at least 1 g is ingested at 0 min to start the ingestion, *i.e*. *u*_0_ ≥ 1. We assumed that ingestion at each time is taken over 0.5 min, and convert *u*_0:60_ to *Glucose* instead of Equation 9, given by

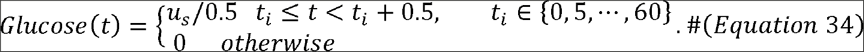

Next, we expressed a nonlinear ordinary differential equation model (Equations 1–8) describing the dynamics of the glucose metabolism system, given by

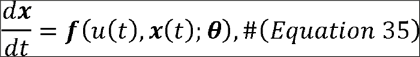

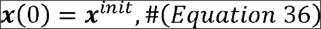

where ***x*** indicates a state variable, ***x***^*init*^ indicates an initial state, ***θ*** is a parameter set, ***f*** and is a nonlinear function. These types and values of ***x***, ***x***^*init*^, **θ**, and ***f*** are different among subjects, because the selected models of subjects and parameters are different among subjects (see STAR Methods B.2). Each subject has one set of ***f***, ***x***^*init*^, and *θ*.***x***(0: *T*). The temporal pattern of ***x*** from *t* = 0 to *t* = *T* with the temporal pattern of oral glucose ingestion ***u***_0:60_ can be obtained by the deterministic numerical simulation of this mathematical model *Sim*, given by

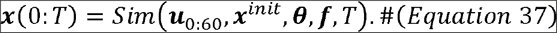

To design a temporal pattern of oral glucose ingestion that minimizes the peak value of blood glucose level, we formulated as an optimization problem. Defining the peak value of blood glucose level in the time course as *x*(0:*T*) as *G*_*Max*_(***x***(0:*T*)) and setting the objective function of the optimization problem to be *J*(*G*_*Max*_(*x*(0:*T*))), we set the objective functions fordesigning the temporal patterns of oral glucose ingestion that minimizes the peak value of blood glucose level, given by

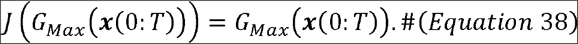

Under these settings, the optimization problem of designing the oral glucose ingestion pattern can be expressed as follows for minimizing the peak value of blood glucose level, given by

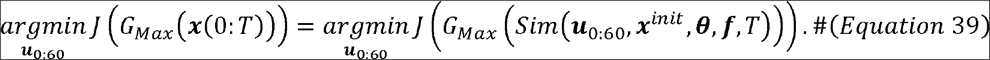

We numerically solved this optimization problem by following evolutionary programming. Each individual has an oral glucose ingestion pattern. After initialization of the oral glucose ingestion pattern of each individual, the algorithm outputs the oral glucose ingestion pattern that minimizes the objective function value by repeating (i) the mutation steps through which a new oral glucose ingestion pattern for each individual is proposed, and (ii) the selection steps through which individual (and thus new pattern) are selected based on the value of the objective function.

Denoting the total number of individuals as *N*, the *n*^th^ individual of the oral glucose ingestion pattern ***u***_0:60_ as ***u***_*n*_, and simplifying the objective function as *J*(***u***_*n*_), the algorithm is as follows:

1. (***Initialization***) For each individual *n* = l,…,*N*, ***u***_*n*_ is initialized and ***u***_*n*_ that minimizes *J*(***u***_*n*_) is stored as ***u***^*^.
2. Repeat the following procedure (a)−(c) *K* times

a. (***Mutation***) For each individual (*n* = 1,…,*N*), copy and mutate ***u***_*n*_ to generate a new individual 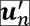. Update ***u***\* as 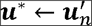 if 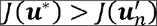.
b. (***Selection* 1**) For each of 2*N* individuals that consist of the original individuals and the new individuals generated at (a), obtain the evaluation value by the following procedure.

i. Select an individual sequentially as ***u***_*m*_.
ii. Select an *M* individuals randomly except for ***u***_*m*_ (duplication possible) as 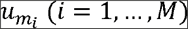.
iii. Obtain the evaluation value defined by the number of 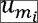 with 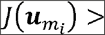 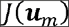
c. (***Selection* 2**) Sort the individuals in order of the evaluation value, and the top *N* individuals are selected and used in the next step.
3. Output ***u***\*.

In terms of evolutionary programming, step 1 is initialization, step 2-a is mutation, and steps 2-b and 2-c are selection. Because the intersection of oral glucose ingestion patterns is complicated by the constraint of 50 g total ingestion dose, this algorithm does not include intersection.

Details of initialization and mutation are as follows: In initialization, to avoid bias of an initial value, *N* individuals consist of an individual with a 50 g bolus ingestion, an individual with 1 g ingestion at 0 min and the remaining 49 g ingestion at 60 min, and other random patterns. The random pattern was generated by distributing 49 g glucose randomly with equal probability at each time point and the remaining 1 g ingestion at 0 min. For the mutation, a new oral glucose ingestion pattern was suggested by repeating operations that transfer 1 g of glucose from one time point to another randomly. Specific operations are as follows.

1. Subtract 1 g of glucose at time 0 min
2. Repeat the following procedure (a) and (b) *L* times

a. Randomly select the source and destination time points of glucose with equal probability.
b. If the ingestion glucose at the source time point contains more than 1 g, transfer 1 g of glucose from the source time point to the destination time point.
3. Add 1 g of glucose at time 0 min

In the deterministic numerical simulation *Sim*, we employed the Euler method with a time step width of 0.001 [min] to shorten the cAICulation time. We also set T = 480 [min].

In the evolutionary programming, we set the number of individuals as *N* = 500, the number of generation except initialization generation as *K* = 500, the number of transfers of glucose in one mutation *L* to decrease from *L* = 20 by 1 every 25 generations, and the number of individuals for cAICulation of evaluation value in selection as *M* = *N*/5 = 100. According to this algorithm and these settings, we cAICulated the optimal ingestion pattern for 5 trials and obtained the pattern that produced the smallest objective function. Note that we obtained the same minimization pattern for each subject multiple times for multiple trials (all trials in subject #1 and #2, 2 trials in subject #3).

#### B.4. Coarse Graining of Minimization Pattern

We coarse-grained the minimization pattern into three periods: a start time (0 min) of the first bolus ingestion, a continuous-like intermediate period, and an end time of the ingestion (Figure 5A, upper panel). The coarse-grained pattern was characterized by 5 parameters: the dose of ingested glucose at 0 min *u*_0_ [g], the start time of the intermediate period *t*_*s*_ [min], the uration of intermediate period *Δt* [min], the dose of ingested of glucose at the end time *u*_*T*_ [g], and the end time of the ingestion *T* [min]. Similar to Figure 4, *t*_*s*_ and *Δt* are multiples of 5 [min]. During the intermediate period (*t*_*s*_ to *t*_*s*_ + *Δt*), the dose of glucose, determined by subtracting *u*_0_ and *u*_*T*_ from 50 g, is equally distributed every 5 minutes. The dose of ngestion during intermittent period is not limited to an integer. These are described as follows:

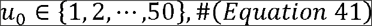

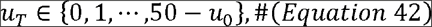

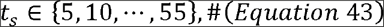

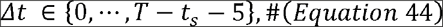

and Equation 22 is replaced by

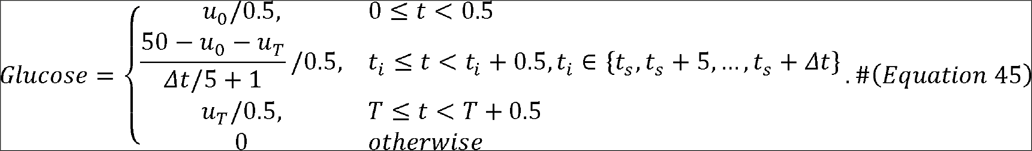

